# A dichotomous family of Cache domain proteins evolved through c-type heme acquisition

**DOI:** 10.64898/2026.02.01.703092

**Authors:** Maithili Deshpande, Jiawei Xing, Vadim M. Gumerov, Robert Dunleavy, Herbert Castillo, Brian R. Crane, Igor B. Zhulin

## Abstract

PAS domains often bind cofactors, whereas related Cache domains are thought to function exclusively as ligand-binding modules. We identify and characterize a large family of Cache domains containing covalently bound c-type heme. Sequence, phylogenetic and structual analysis of the DUF3365/Tll0287 family reveals conservation of a canonical CXXCH heme-binding motif, establishing c-type heme binding as a defining feature of this family. Despite previous annotation as PAS-like, these proteins share a distinct heme-binding insertion into a pre-existing Cache scaffold, representing the first documented recruitment of an intact cofactor-binding motif into a bacterial sensor domain. Domain architecture further divides the family into two subclasses: one exhibiting fusions to signaling domains and another comprised of standalone proteins. Spectroscopic analyses indicate that a representative signaling protein acts as a nitric oxide sensor, whereas a stand-alone representative exhibits redox-related properties and weak nitric oxide binding. These findings expand the functional repertoire of Cache domains and reveal a mechanism for the evolution of bacterial sensory systems.

## Introduction

Signal transduction is a fundamental process that enables cells to perceive and adapt to changing conditions. In bacteria, this process involves a diverse set of sensory proteins that detect a wide range of physical and chemical stimuli, including nutrients, toxins, temperature changes, and signaling molecules^1,2^. Upon sensing these cues, these proteins initiate a cascade of intracellular signals, often through reversible phosphorylation or second messenger production, to modulate gene expression, motility, metabolism, and virulence. The largest families of sensory proteins in bacteria include transcriptional regulators^2,3^, histidine kinases^1^, second messenger-based systems, such as those utilizing cyclic di-GMP^4^ or cAMP^5^, and methyl-accepting chemotaxis proteins (MCPs)^6,7^. Collectively, signaling networks employing these sensory proteins enable bacteria to thrive in diverse environments, contributing to their ecological success.

Sensory proteins allow bacteria to detect various cues both intracellularly and externally. These responses are achieved by having dedicated sensor domains that are fused to downstream signaling modules, such as a histidine kinase catalytic domain and MCP signaling domain. Among the many sensor domains that mediate signal detection, **P**er-**A**rnt-**S**im (PAS) domains are particularly well-studied. PAS domains represent one of the largest superfamilies of intracellular sensory modules that primarily sense redox states, light, and gases, such as oxygen, carbon monoxide (CO), and nitric oxide (NO) by binding cofactors such as flavins or heme^8–10^. By contrast, **Ca**lcium channels and **che**motaxis receptors (Cache) domains that are homologous and structurally similar to PAS domains are found in the extracytoplasmic regions of sensory proteins, where they function as ligand-binding modules^11,12^. Cache domains form the largest family of extracellular sensors in bacteria^12^ and mediate sensing of external small-molecule ligands, such as amino acids, biogenic amines, nucleotides, organic acids, and sugars^13–18^.

In this study, we describe a large family of Cache sensor domains that evolved through acquisition of a short cytochrome c-derived peptide containing a conserved c-type heme–binding motif. The proteins that have these heme-containing Cache domains (designated here as sCache_heme domain) were classified into two main categories: those that appear as standalone domains or those fused to signal transduction modules. To elucidate the possible functions of these proteins, we experimentally characterized two proteins in this family, one from each category. The stand-alone domain from *Pseudomonas putida* shows redox-related properties. The signal transduction protein selected from *Pseudomonas azotoformans* has an unusual architecture with an isolated cytoplasmic HAMP (**h**istidine kinase, **a**denylyl cylclase, **m**ethyl-accepting chemotaxis protein, **p**hoshphatase) domain^19^ and similar kinetic and spectroscopic properties as known nitric oxide sensors. Finally, we show that this sensor is found in many bacterial phyla, with enrichment in Gracilicutes, and it serves as an input to major signal transduction systems in bacteria. Thus, these periplasmic sCache_heme domains perform different functions in context of their heme environment and the domains that they are coupled to.

## RESULTS

### The DUF3365/Tll0287 domain family contains a conserved heme c binding motif

Systematic analysis of bacterial signal transduction proteins in the MiST database^20^ revealed that many sensory proteins, including histidine kinases, MCPs, and c-di-GMP metabolizing enzymes harbor an N-terminal domain of unknown function, originally annotated in the Pfam database^21^ as DUF3365 (PF11845). Following the integration of Pfam into the InterPro resource^22^, this domain was renamed Tll0287-like, retaining the same Pfam accession and assigned InterPro accession IPR021796. This domain family was assigned to the PAS_fold superfamily, following the original interpretation by the authors who solved the structure of the *Thermosynechococcus elongatus* protein Tll0287 (PDB: 5B82) and noted that it adopts a fold resembling PAS sensor domains rather than typical c-type cytochrome scaffolds^23^. They further demonstrated that Tll0287 protein contains *c* heme, which is axially ligated by cysteine and histidine residues and harbors a canonical CXXCH motif that forms thioether bonds with the heme vinyl groups^23,24^.

To comprehensively collect DUF3365/Tll0287 sequences from public databases, we employed two complementary approaches. First, we downloaded a curated set of more than 5,000 protein sequences matching the Tll0287-like domain profile from the InterPro database, which contains curated, taxonomically balanced set of genomes (Dataset S1). To explore a larger sequence space beyond this stringently curated dataset, we performed PSI-BLAST searches against the NCBI Clustered NR database^25^ using the Tll0287 protein sequence as a query, which retrieved a larger set of ∼20,000 sequences (Dataset S2).

In addition to Tll0287, both datasets included two proteins from *Geobacter sulfurreducens*, GSU0582 and GSU0935, which were previously characterized and structurally resolved. These proteins were assigned to the PAS superfamily and shown to contain covalently bound heme c with His/OH_2_(OH^−^) ligation^26^. We constructed multiple sequence alignments of Datasets 1 and 2, which revealed that the canonical CXXCH c-type heme-binding motif is strictly conserved across this domain family. Conservation rates for this motif in Dataset 1 and 2 were 99% and 99.4%, respectively. Together, these results indicate that covalently bound c-type heme is a defining and likely functionally essential feature of the DUF3365/Tll0287 domain family.

### DUF3365/Tll0287 is a cofactor-containing Cache domain

All prior classifications, including original descriptions of the Tll0287^23^ and GSU0582/GSU0935^26^ proteins, as well as Pfam and InterPro annotations, assigned this domain family to the PAS superfamily. Indeed, PAS domains include several families capable of heme-binding^10^. However, we hypothesized that DUF3365/Tll0287 belonged to the Cache rather than PAS superfamily. This hypothesis was motivated by two observations. First, all previously reported “extracytoplasmic PAS domains” defined based on structural interpretations have subsequently been reclassified as Cache domains following more detailed sequence and structure analyses^12^, Second, bona fide PAS domains that bind heme are cytoplasmic and utilize noncovalently bound heme b^27,28^, whereas c-type heme is invariably located in the extracytoplasmic compartment because heme covalent attachment involves membrane transport^29^.

To re-evaluate the family and superfamily assignment of DUF3355/Tll0287, we performed both structure- and sequence-based searches. In structural comparisons, all ten top matches defined by a DALI^30^ search against the PDB database^31^ using the Tll0287 structure (PDB: 5b82) were annotated as Cache domains by InterPro/Pfam (Table S1). Consistently, profile-profile sequence comparisons using HHpred^32^ also assigned DUF3365/Tll0287 to the Cache superfamily (Table S2). The highest-scoring matches included multiple Cache domain family profiles, as well as PDB profiles corresponding to histidine kinases CitA and DcuS and the MCPs PctA and PctC, all of which are well-established members of the Cache superfamily^12,33^.

Structurally, the DUF3365/Tll0287 fold shares a conserved core architecture with ligand-binding Cache domains but contains a distinct structural element that is absent from canonical sCache (comprised of a **<u>s</u>**ingle **<u>Cache</u>** module) domains – an insertion between β3 and β4 containing a short α-helix and adjacent loop regions that encompass the CXXCH motif (Fig. 1). Based on these combined sequence and structural features, we designate this domain **sCache_heme**, representing the first known Cache domain that contains a covalently attached cofactor.

**Figure 1.**
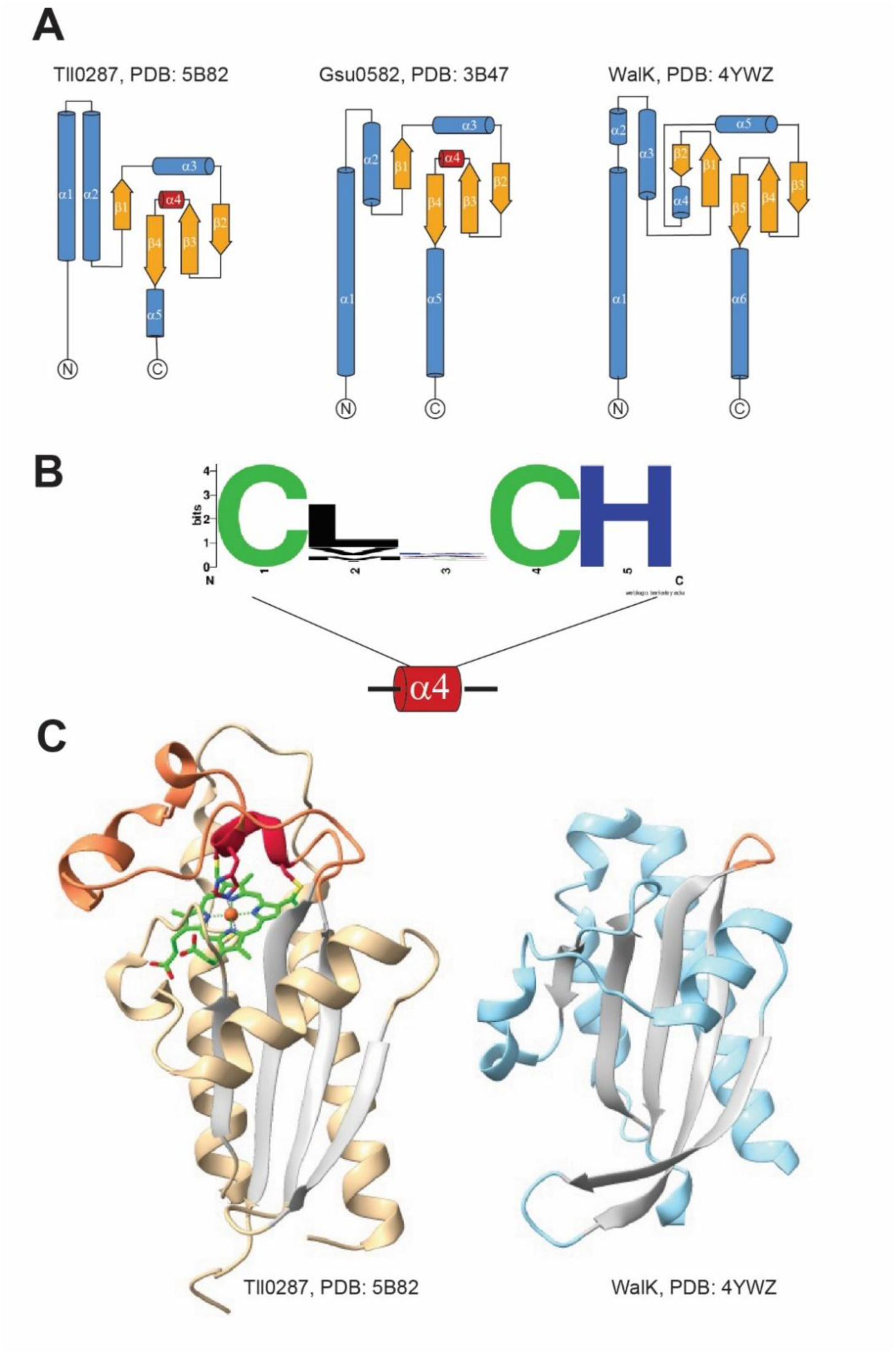
Structural comparison of sCache_heme and sCache domains. **A)** Topology diagrams of periplasmic domains of Tll0287 and Gsu0582 in comparison to the sCache domain of the WalK sensor histidine kinase. Alpha helices are shown in blue and beta strands – in orange. **B**) A short alpha helix (α4) inserted between strands β3 and β4 contains the CXXCH cytochrome c binding site. Sequence logo was generated from a multiple sequence alignment of Dataset 1 (>5,000 sequences). **C**) Side-by-side comparison of Tll0287 and WalK structures. sCache_heme domains have an expanded loop (orange) preceding the last β-strand that contains the heme (green)-binding CXXCH motif (red).

### Phyletic distribution and domain architecture of sCache_heme-containing proteins

sCache_heme domains are distributed across a broad range of bacterial phyla, but are disproportionately enriched within Gracilicutes, particularly in Defferibacterota, Nitrospirota, Desulfobacterota, Campylobacterota, and Aquificota (Table S3, Fig. 2), where more than 50% of sequenced genomes encode at least one sCache_heme-containing protein. In contrast, these proteins are rare or entirely absent from many Terrabacteria phyla, including Bacillota, Actinomycetota, and Chloroflexota. Cyanobacteriota represents the only Terrabacteria with a substantial fraction of genomes (∼30%) encoding sCache_heme (Table S3, Fig. 2). Several deeply branching bacterial phyla – Fusobacteria, Deinococcota, Synergistota, and Thermotogota – are also largely devoid of sCache_heme domains. This phyletic distribution suggests that the sCache_heme emerged within Gracilicutes, followed by limited horizontal transfer into some Terrabacteria lineages.

**Figure 2.**
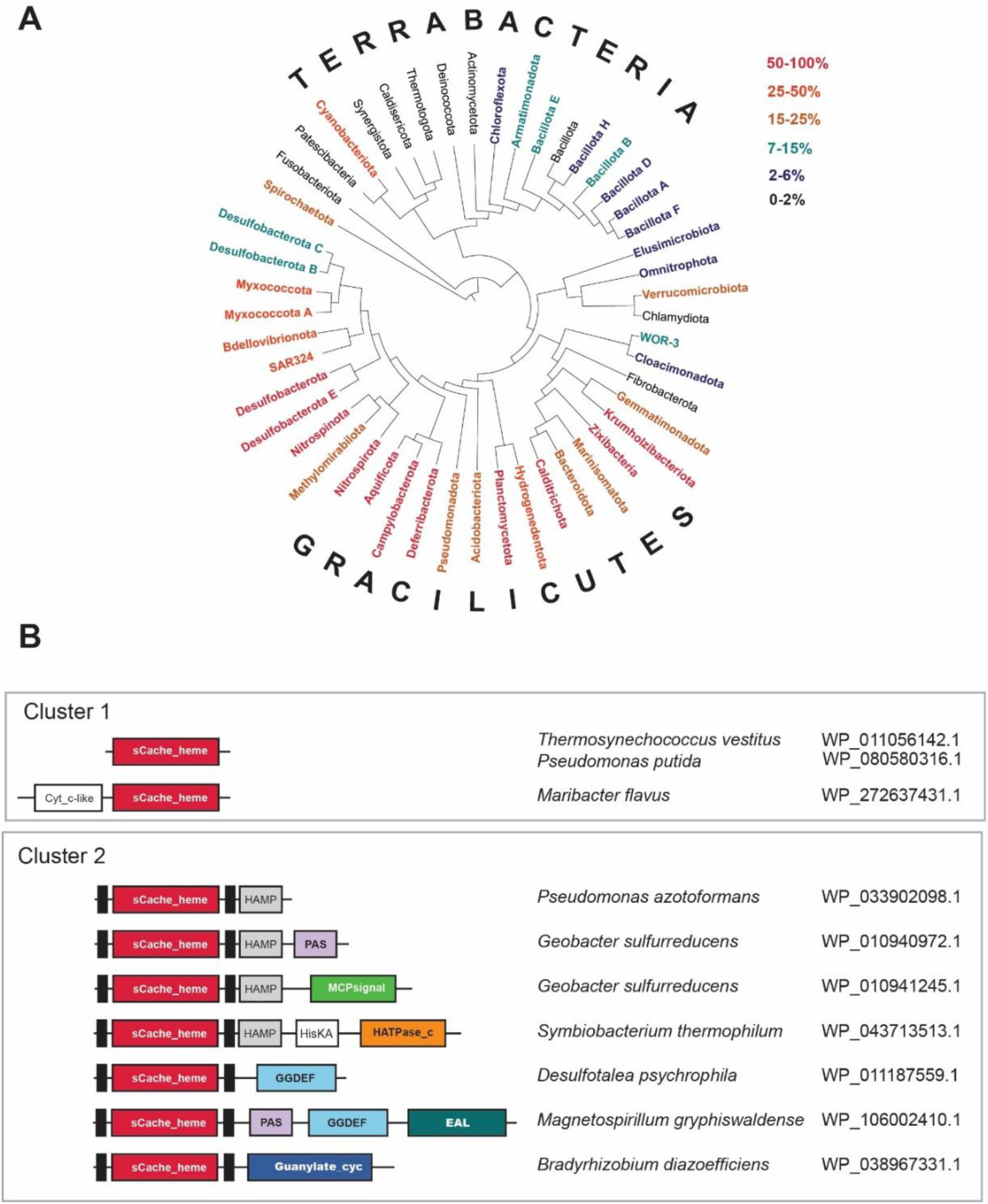
Phyletic distribution and domain architecture of sCache_heme. **A**) GTDB taxonomy tree showing major bacterial phyla. Colors represent the percentage of genomes in a phylum that harbor sCache_heme domain containing proteins. **B**) Most frequent associations of sCache_heme with signaling domains. Representative genomes and accession for sCache_heme proteins are shown. Domain nomenclature according to Pfam: HAMP (PF00672) – present in <u>H</u>istidine kinases, <u>A</u>denylate cyclases, <u>M</u>ethyl-accepting chemotaxis proteins, and <u>P</u>hosphatases; PAS (CL0183) – found in <u>P</u>ER, <u>A</u>RNT, and <u>S</u>IM proteins; MCPsignal (PF00015) – Methyl-accepting chemotaxis protein (<u>MCP</u>) <u>signal</u>ing domain; HisKA (PF00512) – Histidine Kinase phosphor-acceptor domain; HATPase_c (PF02518) – present in Histidine kinase-, DNA gyrase B-, and HSP90-like ATPase; GGDEF (PF00990) – diguanylate cyclase GGDEF domain; EAL (PF00563) - c-di-GMP phosphodiesterase EAL domain; Guanylate_cyc (PF00211) – Adenylate and guanylate cyclase catalytic domain.

Analysis of domain architectures revealed that nearly half of all sCache_heme domains occur as stand-alone domains. This organization is highly unusual for Cache family members, more than 90% of which are typically fused to downstream signaling modules such as histidine kinases, MCPs, or GGDEF/EAL enzymes. Notably, a fusion of sCache_heme with a cytochrome_c-like domain (Fig. 2), which also contains the CXXCH motif, was identified exclusively in Flavobacteria. We designated this group as Cluster 1 (Fig. 2). In the remaining cases, sCache_heme domains are fused to classical signaling outputs consistent with their incorporation into modular signal transduction systems. This group was designated as Cluster 2. This distribution raises the question of whether the domains in Cluster 1 also participate in signaling or instead perform redox-related functions similar to cytochromes.

### Origin of the heme c-binding motif in Cache sensor domains

To explore the likely evolutionary scenario underlying heme acquisition by the sCache_heme domain, we performed two complementary sequence similarity searches. First, we conducted BLAST searches with default parameters against the RefSeq and Clustered NR databases, using a 13–amino acid sequence from GSU0582 (WP_010941245.1; PDB: 3B47) - EVRCQSCHEQGAR - corresponding to α-helix 4 and portions of the adjacent loops. In both searches the highest scoring sequences comprised a diverse set of cytochrome family proteins containing the canonical CXXCH motif (Tables S4 and S5).

Second, we performed PSI-BLAST searches (BLASTP lacked sufficient sensitivity) against the RefSeq database using the sCache domain sequence from which the 13-aa region containing the CXXCH motif had been removed. To ensure generality, we carried out this analysis using multiple sCache_heme domains derived from different protein contexts, including MCPs, histidine kinases, and GGDEF/EAL-containing proteins. In addition to sCache_heme domains, these searches consistently retrieved numerous proteins harboring classical sCache domains that lack the CXXCH motif. As an illustrative example, a PSI-BLAST search initiated with the sCache_heme domain of GSU0582 (with the CXXCH-containing segment deleted) retrieved a DcuS/MalK family sensor histidine kinase from *Priestia aryabhattai* (WP_411982187.1). This sequence was recovered in the sixth iteration with a highly significant E value of 3.28 × 10⁻¹⁷ (Dataset S3). Its N-terminal region (residues 32–166) yielded the top ten Pfam matches to multiple families within the Cache superfamily (Table S6).

Together, these results support a scenario in which a short fragment derived from a cytochrome c–like protein, carrying the heme-binding CXXCH motif, was inserted into a preexisting stand-alone sCache domain. insertion and deletion of sequence fragments are well-established mechanisms of Cache domain evolution. Cache domains vary substantially in length, even among members of the same ligand-binding family, reflecting frequent gain and loss of structural elements during evolution^13–15^. In some cases, such events involve insertion of an entire domain into the middle of the Cache scaffold, as exemplified by the evolution of the eukaryotic amino acid-sensing Cache domain^13^. Thus, inserting a short sequence fragment into a pre-existing Cache domain is fully consistent with the known evolutionary plasticity of this superfamily. The cytochrome-origin model is considerably more parsimonious than alternative scenarios, such as independent evolution of the CXXCH motif within the Cache scaffold. The stepwise emergence of this complex motif would involve intermediate evolutionary states that would be unlikely to support covalent heme attachment or other meaningful function and therefore would be expected to provide no selective advantage.

The stand-alone sCache_heme domains represent the largest group within this protein subfamily, in contrast to ligand-binding Cache domains reported previously^13–15,18^. In this context, fusions of sCache_heme domains with signal transduction modules leading to the emergence of NO-sensing histidine kinases, MCPs, and cyclic nucleotide–modulating enzymes are likely secondary and more recent evolutionary events. Recruitment of enzymatic domains as sensors into existing signal transduction pathways is well documented^34,35^. However, to our knowledge, this represents the first example in which only a small fragment of an enzymatic domain was co-opted into a signaling pathway via insertion into an existing sensor domain.

Finally, only a single archaeal genome in the RefSeq database^36^ - *Methanohalophilus levihalophilus* - encodes a sCache_heme containing sensor histidine kinase (WP_209678719.1). A small number of homologs are detectable in archaeal genomes within the larger NR database^25^ (Dataset S2), consistent with the emergence of this domain after the divergence of bacterial and archaeal lineages and with only exceedingly limited horizontal transfer of sCache_heme domains to archaea.

### Residue conservation within the two clusters of sCache_heme proteins

The two types of sCache_heme proteins also exhibit specific patterns of sequence conservation. Cluster 1 proteins show strong conservation of hydrophobic residues at two positions in α-helix 1: position 44 is occupied by Leu, Val, Ile, or Met in 97% of sequences, and position 48 is occupied by the same set of hydrophobic residues in 95% of sequences. In addition, position 95 within the loop connecting β-strand 1 and α-helix 3 is occupied by Asn or Ser in 93% of Type 1 sequences. These positions are not equivalently conserved in Type 2 proteins.

Conversely, Cluster 2 proteins contain two family-specific conserved charged positions. Position 85 in β-strand 1 is occupied by the positively charged or potentially positively charged residues Arg, Lys, or His in 87% of sequences, whereas position 101 in α-helix 3 is occupied by the negatively charged residues Asp or Glu in 86% of sequences. Thus, Cluster 1 is distinguished primarily by conservation of hydrophobic residues in α-helix 1 and a polar residue in the adjacent loop region, whereas Cluster 2 is characterized by conservation of oppositely charged residues in β-strand 1 and α-helix 3. Although the functional significance of these family-specific positions remains unknown, their conservation indicates distinct sequence specialization within the two architectural classes.

### Target selection for experimental validation

The evolutionary scenario proposed above predicts that sCache_heme domains associated with signal transduction modules (Cluster 2) function as heme-based sensors, whereas stand-alone or cytochrome-associated (Cluster 1) may instead serve redox-related roles. To investigate if proteins from each Cluster indeed have different properties, we selected two representatives from the phylum *Pseudomonadota*: a stand-alone sCache_heme protein from *Pseudomonas putida* designated as Target 1 (Cluster 1, Fig. 2) and a membrane-associated sCache_heme protein from *P. azotoformans* containing a transmembrane domain and a cytoplasmic HAMP domain designated as Target 2 (Cluster 2, Fig. S1). Target 1 is closely related to the canonical Tll0287 protein and was predicted to coordinate heme via Cys/His ligation and exhibit properties consistent with redox functions. Due to the presence of a HAMP domain, which has strictly signaling properties^19,37^, Target 2 was predicted to form a signaling-competent dimer with a more labile heme coordination environment.

### Expression, purification, and heme c incorporation

Both sCache_heme proteins were heterologously produced in *E. coli* (Fig. S1) by co-expression with the cytochrome c (Cc) maturation (Ccm) system on plasmid pEC86, which provides constitutive expression of eight Ccm proteins under aerobic conditions. The Ccm system covalently attaches c-type hemes in the periplasm of gammaproteobacteria (including *Pseudomonas* and *E. coli* spp.) and is required for proper protein folding.

The Target 1 protein from *P. putida* was purified as a maltose binding protein (MBP)-fusion as MBP both greatly improved protein solubility and also transported the protein to the periplasm for heme c incorporation. The *P. putida* sCache protein forms inclusion bodies when expressed with the native *Pseudomonas* signal peptide or the *E. coli* PelB signal peptide. MBP could be removed by protease cleavage without change of the protein spectral properties, but was usually retained to improve yields. The full-length Target 2 membrane protein from *P. azotoformans* was solubilized in the detergent n-dodecyl-β-D-maltoside (DDM) and purified by Ni-NTA affinity chromatography, followed by desalt and size-exclusion chromatography (SEC). SEC-SAXS analysis indicated that the *P. azotoformans* protein forms a stable dimer in detergent solution (Fig. S3). Both proteins were highly pure (>90% purity) after SEC (Fig. 3, Figs. S2 and S3) and had a high proportion of heme incorporation (Fig. S4).

**Figure 3.**
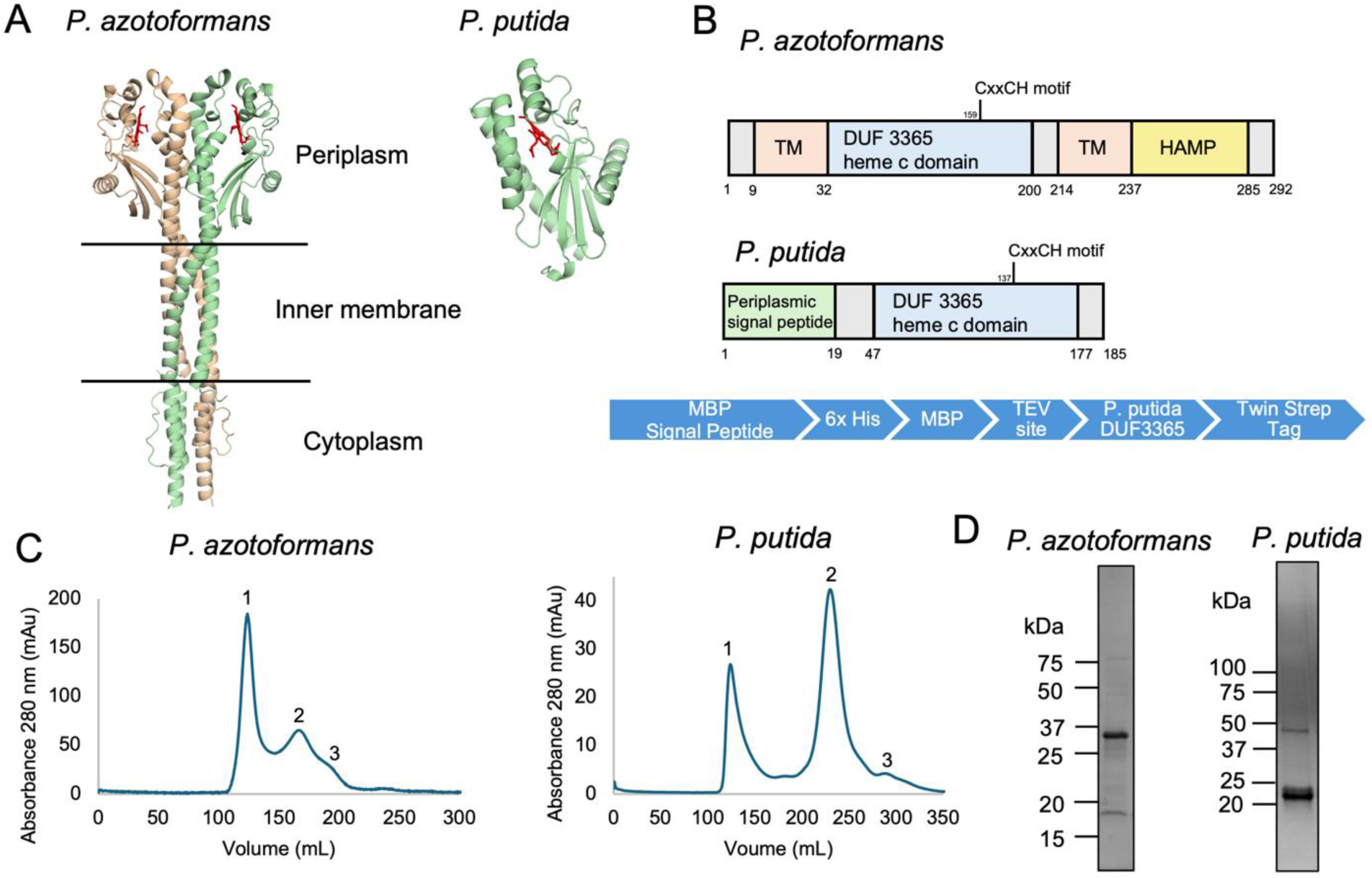
**Purification of the sCache_heme domain proteins from *P. azotoformans* and *P. putida.*** A. AlphaFold 3 models of the *P. azotoformans* and *P. putida* sCache_heme domain containing proteins. The c-type heme in the periplasmic sCache_heme domain is colored red. The signal peptide for the *P. putida* protein is cleaved during transport to the periplasm and is not present in the structure. B. Domain organization of the sCache_heme proteins. The *P. putida* sCache_heme protein was expressed as an MBP-fusion with a tobacco-etch virus (TEV) protease cleavage site and the construct with purification tags is shown in blue. C. Size-exclusion chromatography elution profiles of the sCache_heme proteins on an S200 26/60 column. For the *P. azotoformans* protein, peaks 2 and 3 were pooled for further experiments. The MBP-*P. putida* fusion protein was cleaved overnight with TEV protease in the absence of reducing agents, as they affect the heme moiety. The cleaved protein elutes at peak 2. D. Coomassie-stained SDS-PAGE gel of purified *P. azotoformans* and *P. putida* sCache_heme proteins (34.5 kDa (monomer) and 21.15 kDa, respectively).

### Spectroscopic characterization of redox states

UV-visible absorption spectra of both proteins revealed features characteristic of low-spin six-coordinate c-type heme in reduced and oxidized states. (Fig. 4, Table 1). Target 2 lacks the Cys residue in the distal heme pocket of Target 1 and no protein-derived ligand is predicted to be in the vicinity of the axial coordination position. The Target 2 spectra for both the oxidized and reduced states resemble those of the Met80Ala iso-1-cytochrome c^38^, which coordinates hydroxide under alkaline conditions, such as those used here to stabilize the protein (pH 8.5). The spectrum of the *P. putida* Target 1 protein is unaltered by the MBP tag (Fig. S5) and is very similar to that of Target 2 after one round of reduction and re-oxidation at pH 8.5 (Figs. 4, S6), yet slightly shifted compared to a variant with a Cys-to-Ala substitution (C59A) in the residue analogous to the heme ligand of Tl0287 (Figs. S6-7). In both the WT and C59A variant we observed a characteristic blue shift of the Soret band and flattening of the β-band as pH is lowered, which is representative of His/OH^−^ ligation (Fig. S8). However, compared to these hydroxide-bound forms, the “as purified” spectra of the *P. putida* protein has a blue-shifted Soret band at pH 8.5, a more pronounced β-band, and a more intense 354 nm band, similar to a variant of iso-1-cytochrome c with Cys/His ligation^39^ (Fig.4, S6). Thus, the Target 1 protein as purified, may be at least partially Cys-ligated, like Tll0287, but upon reduction converts to His/OH^−^ ligation. Indeed, reduction potentials of the WT *P. putida* protein (123 mV) and the C59A variant (113 mV) at pH 7.0 (Table 2, Figs. S9-S10) are inconsistent with Cys/His ligation, which are usually much lower (∼ -200 to - 400 mV)^39,40^. Although reduction potentials of heme proteins span large ranges across diverse functions, the *P. putida* potential does fall within the range of c-type cytochromes that act as periplasmic electron carries in various metabolic pathways and respiratory chains^41^ (e.g. c_551-3_, c_4_) and also aligns with that of the periplasmic nitric oxide carrier, cytochrome c’^42^.

**Figure 4.**
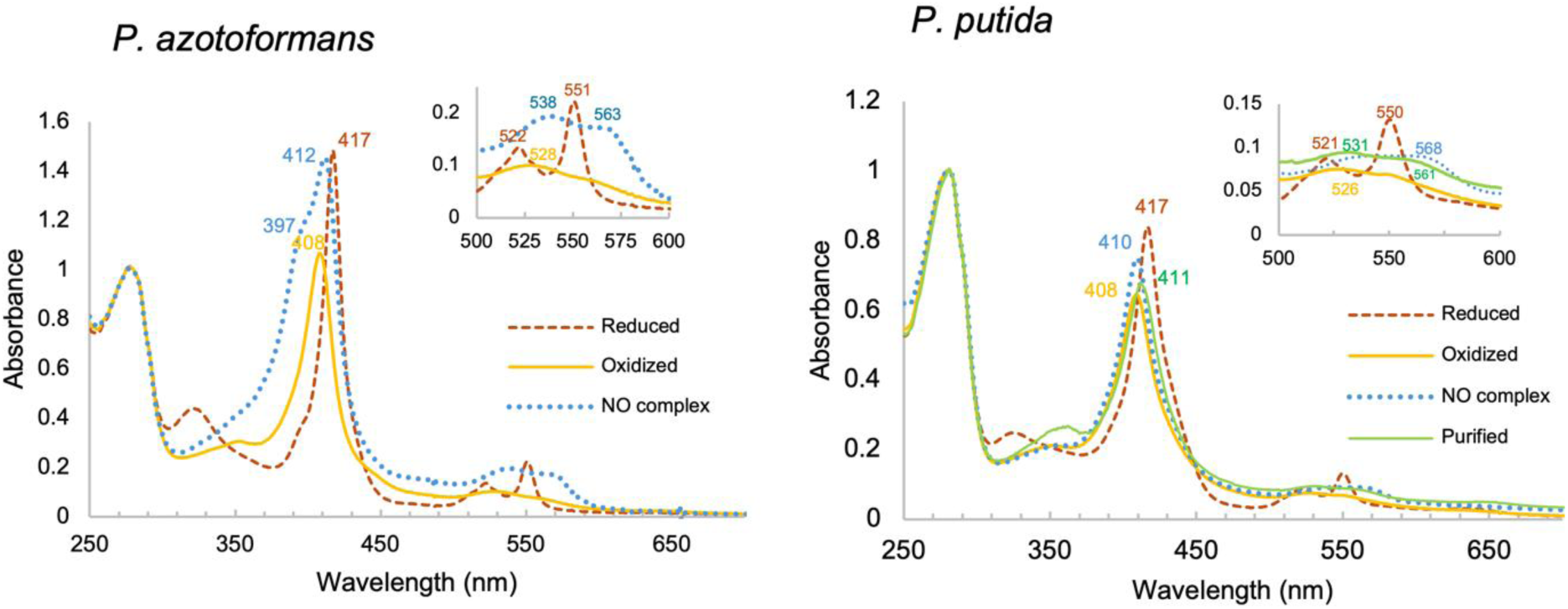
UV-Vis absorption spectra of the *P. azotoformans* and *P. putida* sCache_heme proteins in oxidized (yellow ━━), reduced (red -----), Fe^2+^-NO forms (blue ^…….^) and protein as purified (green ━━). The insert shows an expanded view of the α and β peaks. Both proteins, on anaerobic reduction with dithionite followed by oxidation in air, gave Soret maxima at 417 nm with split α and β bands at ∼550 nm and ∼521 nm in the Fe²⁺ (reduced) state and Soret maxima at 408 nm with a broad α/β band at ∼526 nm in Fe³⁺ (oxidized) state at pH 8.5. The Fe^2+^-NO spectra for *P. putida* sCache_heme showed Soret maxima at 410 nm and relatively flat, diffused α/β peaks with features at ∼568 nm and ∼530 nm. In contrast, the *P. azotoformans* protein gave spectra with a 412 nm Soret peak and 397 nm shoulder, and distinct α and β bands. The *P. putida* sCache_heme protein upon purification had Soret maxima at 411 nm, distinct from that obtained on reduction and subsequent oxidation of the protein.

**Table 1.** Soret, α and β peaks of the sCache_heme proteins, c-type heme proteins, and NO sensors.

| | Soret<br>(nm) | $\beta$<br>(nm) | $\alpha$<br>(nm) | Reference |
| --- | --- | --- | --- | --- |
| **Fe<sup>2+</sup>** |  |  |  |  |
| *P. azotoformans* sCache_heme | 417 | 522 | 551 | this work |
| *P. putida* sCache_heme | 417 | 521 | 550 | this work |
| Cytochrome c (pH 6.8) | 415 | 521 | 550 | 66 |
| *Rc* cytochrome c' (RCCP) | 425/435 | 550 |  | 45 |
| GSU0582 | 415 | 522 | 551 | 26 |
| GSU0935 | 415 | 522 | 550 | 26 |
| sGC | 431 | 555 | 555 | 67 |
| *Pa* NosP | 420 | 524 | 554 | 52 |
| *Tt* H-NOX | 431 | 565 |  | 68 |
| **Fe<sup>3+</sup>** |  |  |  |  |
| *P. azotoformans* sCache_heme | 408 | 528 |  | this work |
| *P. putida* sCache_heme | 408 | 526 |  | this work |
| Cytochrome c (pH 6.8) | 409 | 530 |  | 66 |
| GSU0582 (physiological pH) | 401 |  |  | 26 |
| GSU0582 (alkaline pH) | 410 |  |  | 26 |
| GSU0935 (alkaline pH) | 408 |  |  | 26 |
| **Fe<sup>2+</sup>-NO** |  |  |  |  |
| *P. azotoformans* sCache_heme | 397/412 | 538 | 563 | this work |
| *P. putida* sCache_heme | 410 | 560 (diffused) |  | this work |
| Cytochrome c (pH 13) | 411 | 538 | 567 |  |
| *Rc* cytochrome c' (RCCP) | 395/414 | ~540 | ~570 | 39 |
| *Ax* cytochrome c' (AXCP) | 415 | 540 | 575 | 38 |
| sGC | 398 | 537 | 571 | 58 |
| *Pa* NosP | 396 | 534 | 574 | 56 |
| *Tt* H-NOX | 420 | 547 | 575 | 68 |
| **Fe<sup>2+</sup>-CO** |  |  |  |  |
| *P. azotoformans* sCache_heme | 415 | 533 | 560 | this work |
| *P. putida* sCache_heme | 413 | 529 | 550 | this work |
| Cytochrome c (pH 14) | 415 | 533 | 560 | 66 |
| *Rc* cytochrome c' (RCCP) | 417 | 535 | 566 | 45 |
| sGC | 423 | 541 | 567 | 67 |
| *Pa* NosP | 416 | 538 | 565 | 52 |
| *Tt* H-NOX | 424 | 544 | 565 | 68 |

**Table 2.** *P. putida* sCache reduction potentials and comparisons*.

| c-type Cytochrome | Reduction potential (mV) | Reference |
| --- | --- | --- |
| *P. putida* sCache_heme WT | 123 ± 4 | This work |
| *P. putida* sCache_heme C59A | 113 ± 1 | This work |
| Cytochrome c (horse heart) | 260 | 69 |
| *Pseudomonas aeruginosa* cytochrome c-551 | 250 | 70 |
| *Neisseria gonorrhoeae* cytochrome c-4 (B) | 217 | 71 |
| *Chlorobaculum thiosulfatophilum* c<sub>555</sub> | 145 | 72 |
| *Nitrosomonas europaea* cytochrome c<sub>co</sub>-550 | 140 | 73 |
| *Alicyclophilus denitrificans* cytochrome c' | 132 | 74 |
| *Sinorhizobium meliloti* SorU | 110 | 75 |
| *Rhodopseudomonas palustris* cytochrome c' | 102 | 74 |
| *Nitrosomonas europaea* cytochrome c-554 (A,B) | 50, 32 | 76 |
| *Desulfovibrio vulgaris* Hildenborough c<sub>553</sub> | 20 | 77 |
| *Nitrosomonas europaea* cytochrome c<sub>co</sub>-552 | -50 | 73 |

\*Midpoint reduction potentials (E°) of protein-bound heme cofactors were determined by spectrophotometric redox titrations using 5-deazariboflavin-mediated photoreduction and 1,2- naphthoquinone as the reference redox couple. Measurements were performed at pH 7.0 and 10 °C. Values represent the mean ± standard deviation vs SHE from 3 independent experiments. Redox potentials of other cytochromes are listed for comparison.

The absorption spectra of the two proteins were strikingly different in the Fe^2+^-NO states. The *P. putida* protein formed a six-coordinate Fe^2+^-NO complex with spectral features resembling that of canonical cytochrome c (Table 1), whereas the *P. azotoformans* protein formed a mixture of five- and six-coordinate Fe²⁺–NO species (Fig. 4).

### NO binding and oxygen reactivity distinguishes sensory and redox-like sCache_heme proteins

Despite similar Fe^3+^ and Fe^2+^ spectra, the two proteins differed markedly in their response to nitric oxide and oxygen. The *P. putida* WT and C59A variants exhibited rapid NO dissociation, indicating weak NO binding (Fig. 5). In contrast, the *P. azotoformans* protein displayed slow NO dissociation, with a k_off_ of (6.25 ± 0.64) × 10⁻⁴ s⁻¹, comparable to established bacterial NO sensors such as NosP and H-NOX (Table 3). Although k_off_ alone does not reflect affinity for NO, it is usually the determining factor as k_on_ values typically vary less than an order of magnitude for NO responsive proteins^43^. NO-induced trans axial ligand dissociation is often a feature of signaling mechanisms by H-NOX proteins. The mixture of five- and six-coordinate NO species formed by the *P. azotoformans* protein has been observed in some cytochrome c’ proteins from photosynthetic bacteria^44–46^, and in NO sensor *Nostoc punctiforme* (*Np*) H-NOX^47^. Unlike *Nostoc punctiforme* H-NOX, where the equilibrium between the two ligation states has a distinct temperature dependence, the mixture of 5- and 6-coordinate species persists in the *P. azotoformans* protein at temperatures of 4, 20 and 40 °C (Fig. S11).

**Figure 5.**
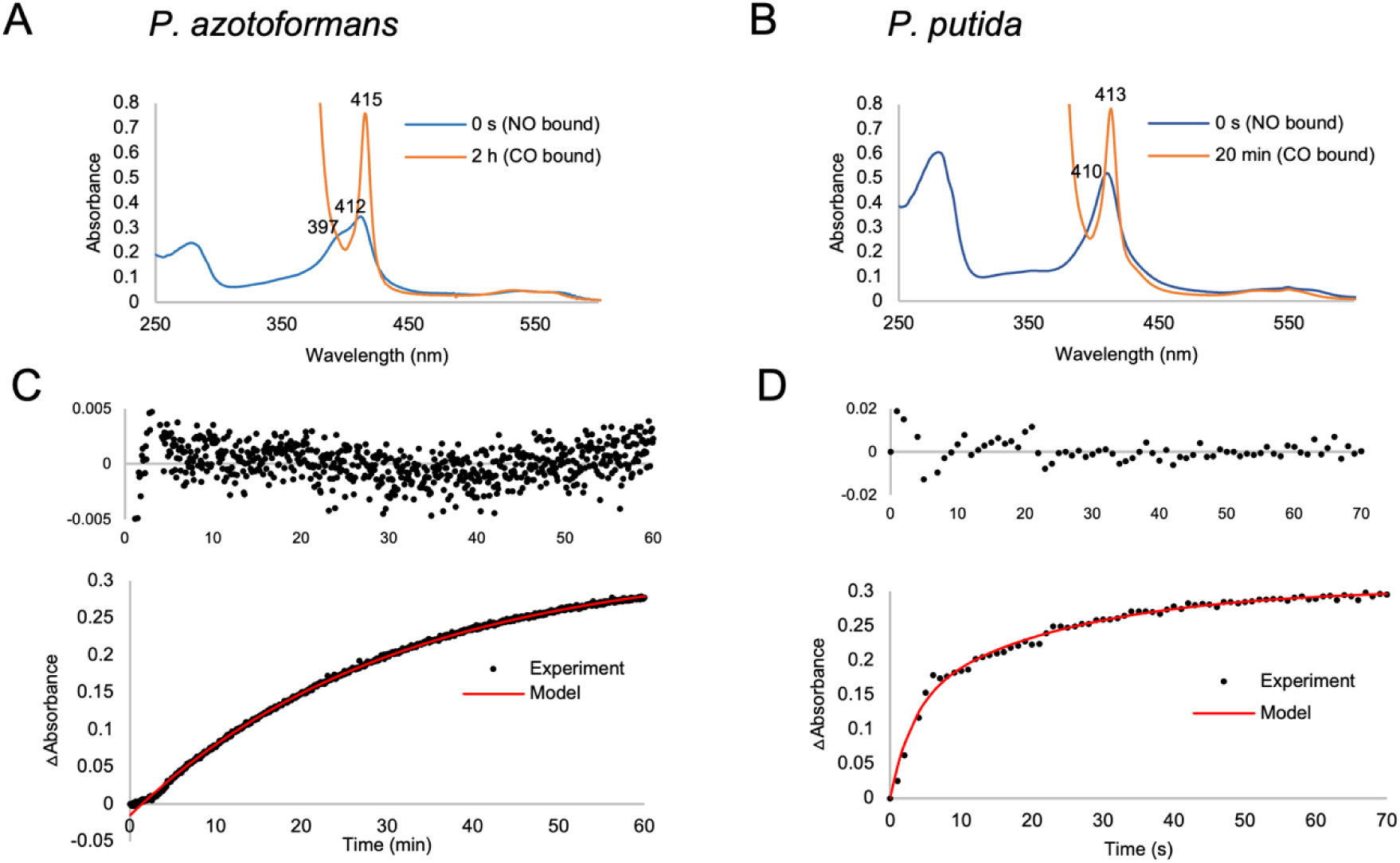
Dissociation of NO from the *P. azotoformans* and *P. putida* Fe^2+^ sCache_heme proteins. A, B. The UV-Vis absorption spectra show conversion of the initial Fe^2+^-NO species (blue) to the final Fe^2+^-CO trap species (orange) upon addition of CO-saturated sodium dithionite buffer into the anaerobic protein solution with a gas-tight syringe. C, D. The increase in absorbance of (C) *P. azotoformans* and (D) *P. putida* CO-bound species was fit to an exponential equation. NO has high affinity for the *P. azotoformans* sCache_heme protein and dissociates slowly over the timescale of one hour, whereas NO dissociates rapidly from the *P. putida* protein within minutes.

**Table 3.** NO dissociation rate constants.

| Protein | $k_1$ (s<sup>-1</sup>) | $k_2$ (s<sup>-1</sup>) | Reference |
| --- | --- | --- | --- |
| *P. azotoformans* sCache | $(6.25 \pm 0.64) \times 10^{-4}$ | NA | this work |
| *P. putida* sCache | $(6.11 \pm 1.54) \times 10^{-2}$ | $(5.73 \pm 2.86) \times 10^{-1}$ | this work |
| sGC | $3.6 \times 10^{-4}$ | NA | 67 |
| SwH-NOX | $15.2 \times 10^{-4}$ | NA | 78 |
| NosP | $1.8 \times 10^{-4}$ | NA | 52 |
| Cytochrome *bd* | $1.6 \times 10^{-3}$ | NA | 79 |
| Cytochrome *bo* | $3 \times 10^{-2}$ | NA | 79 |
| Cytochrome c oxidase *aa3* | $1.0 \times 10^{-2}$ | NA | 80 |
| Cytochrome *cd1* | $4.35 \times 10^1$ | NA | 81 |
| Cytochrome c554 | $3.4 \times 10^{-2}$ | NA | 82 |
| Cytochrome P450<sub>CAM</sub> | $3.5 \times 10^{-1}$ | NA | 83 |

The reduced *P. azotoformans* protein also autoxidizes ∼10x faster than the reduced *P. putida* protein (k = (4.78 ± 0.83) x 10^-2^ s^-1^ compared to k = (0.5 ± 0.1) x 10^-2^ s^-1^). This likely reflects a more accessible distal heme pocket and coordination site that heighten oxygen reactivity (Fig. S12). Interestingly, the C59A *P. putida* substitution does not greatly affect autoxidation rates, thereby suggesting that both this variant and the WT are likely hydroxide ligated at pH 8.5 (Fig. S13). Regardless, none of the *P. putida* WT, *P. putida* C59A or *P. azotoformans* proteins form a stable complex with oxygen, as the *P. azotoformas* protein does with NO (Fig S12). All readily autoxidize orders of magnitude faster than known O_2_–binding proteins^48^, producing superoxide as a product (Fig. S14). Together, these results demonstrate that sCache_heme domains associated with signal transduction architectures have some properties consistent with nitric oxide sensors (an exchangeable ligand coordination site, low O_2_ binding, slow NO-off rates, and a covalently coupled signal transduction domain), whereas stand-alone sCache_heme domains exhibit properties more consistent with redox-related roles (an axial protein ligand at physiological pH, low O_2_ binding, low NO binding, a typical cytochrome redox potential and no coupled domains).

## DISCUSSION

In this study, we identify and characterize a previously unrecognized class of Cache sensor domains that bind covalently attached c-type heme and perform distinct biological functions depending on their domain architecture. Through comprehensive sequence analysis, we demonstrate that the DUF3365/Tll0287 family universally contains a highly conserved CXXCH motif, establishing c-type heme binding as a defining and ancestral feature of this domain family. Structure- and sequence profile-based comparisons further resolve earlier ambiguities in domain classification by demonstrating that these proteins belong to the Cache superfamily rather than PAS, underscoring the importance of integrating subcellular localization, cofactor chemistry, and evolutionary context when assigning sensor domains.

Phylogenetic and architectural analyses reveal that sCache_heme domains are broadly distributed across bacterial phyla but show striking differences in domain organization. Approximately half occur as stand-alone domains or in association with cytochrome c–like proteins, whereas the remainder are fused to canonical signal transduction modules such as histidine kinases, MCPs, or cyclic nucleotide enzymes. This dichotomy suggested functional divergence within the family, which we tested experimentally by characterizing representative members from each architectural class.

Biochemical and spectroscopic analyses of representative sCache_heme proteins from *Pseudomonas putida* (stand-alone) and *P. azotoformans* (signaling-associated) revealed that both proteins bind c-type heme and form low-spin species in reduced and oxidized states at high pH. However, despite these similarities, their ligand-binding behaviors differ markedly. The *P. putida* sCache_heme protein, which closely resembles the single-domain protein Tll0287, may exhibit some degree of Cys/His ligation in its purified form, but upon reduction, converts predominantly to a His/OH^−^ ligation state. Indeed, this protein binds neither oxygen nor nitric oxide strongly, has a reduction potential typical of redox-active cytochromes (123 mV) and may instead play a role in electron transfer or redox sensing. Although this reduction potential also matches that of cytochrome c’, unlike cytochrome c’, the protein does not stabily bind nitric oxide. We do note that the redox and oxygen-binding parameters of recombinantly expressed and purified heme proteins can be misleading for assigning sensing functions ^49,50^; however, in addition to their different biochemical properties, the stand-alone sCache_heme proteins have different gene architectures and sequence conservation patterns than the signaling-associated proteins. Even stand-alone sensors, such as H-NOX and NosP genetically associate with signaling output proteins like sensor kinases^51,52^. The stand-alone sCache_heme genes show no such patterns – a fact that will make delineation of their true functions an interesting direction to pursue, but also a challenging one.

In contrast, the *P. azotoformans* sCache_heme protein, which is associated with transmembrane helices and a cytoplasmic signal-transmitting HAMP domain, displays properties characteristic of sensors. This protein binds nitric oxide with low off rates while failing to form a stable oxy-complex. Its NO dissociation kinetics are comparable to those of established bacterial NO sensors such as NosP and H-NOX, despite its extracytoplasmic localization and covalently bound c-type heme. The formation of mixed five- and six-coordinate Fe²⁺–NO species, together with rapid autoxidation in air, suggests a more labile heme coordination environment that facilitates ligand binding and signal initiation. We have recently found that a homolog of the *P. azotoformans* interacts directly with a Ser/Thr kinase and that NO binding the periplasmic sCache_heme domain alters the phosphorylation of the “receptor” by the kinase, thereby indicating that NO binding in the periplasm transmits a conformational signal to the cytoplasmic HAMP domain – a property consistent with signal transduction^53^.

Few periplasmic proteins are known to selectively bind NO. Group 1 proteobacterial cytochrome c’ is a rare example of a periplasmic NO binding protein with defined function: NO scavenging for protection against nitrosative stress^54,55^. The Group 1 cytochromes c’ from *Rhodobacter capsulatus* Cc’ (RCCP) has a solvent accessible channel to the distal heme, like the *P. azotoformans* protein, whereas the related group 2 enzymes, such as that from *Alcaligenes xylosoxidans* (AXCP), do not^45^. Interestingly, RCCP forms a mixture of five-and six-coordinate ferrous nitrosyl species with UV-Vis spectra, also very similar to those of the *P. azotoformans* dCache protein, whereas AXCP forms a six-coordinate nitrosyl species like the *P. putida* protein. Thus, the Cluster 2 sCache_heme family members share additional features of NO-reactive proteins.

These findings suggest a functional division within the sCache_heme family: stand-alone domains exhibit properties consistent with redox-related roles, whereas domains fused to signal transduction modules can stably react with exogenous ligands, such as NO. This architecture– function relationship parallels, at a conceptual level, the distinction observed among cytoplasmic heme b–binding PAS domains, while revealing an analogous sensing strategy operating in the extracytoplasmic compartment using c-type heme.

From an evolutionary perspective, our results support a model in which a short cytochrome c– derived fragment containing the heme-binding motif was inserted into a preexisting Cache scaffold, giving rise to sCache_heme. Stand-alone sCache_heme domains likely represent the ancestral state, with subsequent fusion to transmembrane and cytoplasmic signaling modules enabling their recruitment into signal transduction pathways. To our knowledge, this is the first documented example of such fragment-level recruitment of a cofactor-binding motif during the evolution of a bacterial sensor domain.

Together, these findings expand the functional repertoire of Cache domains to include covalently bound heme. A subset of these proteins has characteristics of sensors that may allow them to respond to host-derived NO, microbially produced NO, or endogenous nitrosative stress, depending on ecological context.

## Materials and Methods

### Bioinformatics tools and databases

MiST 4.0 database^20^ was used for scanning bacterial signal transduction proteins for putative novel sensor domains. The InterPro database^22^ was used to download sequences matching the Tll0287-like domain profile. BLAST and PSI-BLAST searches^56^ against the NCBI Clustered NR^25^ and RefSeq^36^ databases were performed with default parameters. The Pfam database^21^ and the TREND webserver^57^ were used for assessment of protein domain architecture. Structure-based searches against the PDB database^31^ were performed using the DALI server^30^ and profile-profile sequence similarity searches were carried out using HHpred^32^. GTDB (Genome Taxonomy Database) release 10^58^ was used to retrieve taxonomy information and the genome-based, phylum-level phylogenetic tree, which was edited and visualized using iTOL v.6^59^. Multiple sequence alignments were constructed using the L-INS-i algorithm of the MAFFT package (v. 7.490)^60^ and visualized and edited in Jalview v. 2.11.3.064^61^. Sequence logo was generated using WebLogo 3^62^. The AlphaFold Protein Structure Database (AFDB)^63^ was used to retrieve structural models.

### Cloning

Codon-optimized full-length sCache_heme domain-containing proteins from *Pseudomonas putida* and *Pseudomonas azotoformans* (accession numbers WP_080580316 and WP_033902098, respectively) were synthesized by Biomatik (Cambridge, Ontario, Canada) custom gene synthesis service. The *P. putida* DUF 3365 protein lacking its native periplasmic signal-peptide was cloned into pMAL-p2G as an MBP-fusion protein with an N-terminal 6x His tag, TEV protease cleavage site and C-terminal twin-Strep tag using kinase, ligase and Dpn1 (KLD) cloning. The *P. azotoformans* DUF 3365 protein was cloned into pET28a with an N-terminal 6x His tag between restriction enzyme sites Nde1 and Xho1 using Gibson assembly. The *P. putida* C59A variant protein was generated by QuikChange site-directed mutagenesis. All plasmids were confirmed by DNA sequencing at the Cornell University Biotechnology Center.

### Protein expression and purification

#### Purification of the P. azotoformans sCache_heme protein

The *P. azotoformans* FL DUF 3365 gene in pET28a was co-expressed with plasmid pEC86 in *E. coli* BL21 (DE3). pEC86 encodes eight cytochrome c (Cc) maturation proteins that are required for the transport and covalent attachment of heme to apo c-type heme proteins^64^. Cells were grown in Terrific broth (IBI Scientific) containing 25 µg/mL chloramphenicol and 50 µg/mL kanamycin at 37°C to an OD 600 of ∼0.8 and induced with 0.5 mM IPTG at 18°C overnight. Cultures were supplemented with 25 µg/mL 5-aminolevulinic acid (Chem-Impex International) at an OD 600 of ∼0.4 to increase heme synthesis. Cells were harvested after ∼16 hr by centrifugation at 4000 x g for 15 min at 4 °C.

To isolate membranes, cells were resuspended in lysis buffer (25 mM Tris pH 8.5, 500 mM NaCl, 10% glycerol), incubated with 0.25 mg/mL lysozyme and lysed using an Avanti Emulsiflex C3 high pressure homogenizer. Membrane fractions were prepared by resuspending the pellet in lysis buffer after ultra-centrifugation of the lysate at 150,000 x g. Protein was solubilized by incubating membranes in lysis buffer supplemented with 1% n-dodecyl-β-D-maltoside (DDM) (Chem-Impex International) for 90 min at 4°C. Insoluble material was pelleted by centrifugation at 75,000 x g for 1 hr. The clarified supernatant was incubated with Ni-NTA resin for 1 hr and washed with 20 column volumes of wash buffer (25 mM Tris pH 8.5, 500 mM NaCl, 10% glycerol, 30 mM imidazole, 0.1% DDM). Protein was eluted in elution buffer (25 mM Tris pH 8.5, 500 mM NaCl, 10% glycerol, 300 mM imidazole, 0.1% DDM) and then quickly desalted into desalt/GF buffer (25 mM Tris pH 8.5, 250 mM NaCl, 5% glycerol, 0.03% DDM) to remove imidazole. The desalted protein was then run over an S200 26/60 gel filtration column pre-equilibrated with desalt/GF buffer. The protein was stored at 4°C and used within 72 hr.

#### Purification of the P. putida sCache_heme protein

The MBP-*P. putida* DUF 3365 fusion protein in pMAL-p2G was co-expressed with plasmid pEC86 in *E. coli* BL21 (DE3). Cells were grown in Luria-Bertani broth (Difco) containing 25 µg/mL chloramphenicol and 100 µg/mL ampicillin to an OD 600 of ∼0.3. Cultures were then supplemented with 25 µg/mL 5-aminolevulinic acid and induced with 0.5 mM IPTG at an OD 600 of ∼0.6. Cells were harvested after ∼16 hr by centrifugation at 4000 x g for 15 min at 4°C. Cells were resuspended in 25 mM Tris pH 8.5, 500 mM NaCl, 10% glycerol and lysed by homogenization. Lysate was centrifuged at 75,000 x g for 1 hr and clarified lysate was incubated with Strep-Tactin XT 4Flow resin (IBA Lifesciences) for 1 hr at 4°C. Resin was washed with 15 column volumes of lysis buffer and protein was eluted in 25 mM Tris pH 8.5, 500 mM NaCl, 10% glycerol, 50 mM D-biotin. The protein was then run over an S200 26/60 gel filtration column pre-equilibrated with GF buffer (25 mM Tris pH 8.5, 250 mM NaCl, 5% glycerol). Fractions containing protein were concentrated, flash-frozen in liquid nitrogen and stored at -80°C.

For cleavage with TEV protease, the MBP-fusion protein was washed and eluted into lower salt buffer (25 mM Tris pH 8.5, 150 mM NaCl, 10% glycerol, 50 mM D-biotin) from Strep-Tactin XT 4Flow resin. Protein was cleaved overnight with a high amount of TEV protease (1:5 molar ratio of MBP-*P. putida* protein to TEV protease) in the absence of commonly used reducing agents DTT or BME required to prevent disulfide bond formation for TEV protease activity. These reducing agents were found to permanently affect the stability and properties of the c-type heme during protein purification, even on removal, and thus could not be used. Protein was then run over Ni-NTA resin to remove TEV protease and uncleaved protein, before proceeding to size-exclusion chromatography as described above. Fractions containing protein were concentrated, flash-frozen in liquid nitrogen and stored at -80°C.

### Size-exclusion chromatography coupled small-angle x-ray scattering (SEC-SAXS)

SEC-SAXS experiments on the *P. azotoformans* sCache_heme protein were performed at the Cornell High Energy Synchrotron Source (CHESS). Protein sample at a concentration of 3 mg/mL was loaded onto a Superdex 200 5/150 GL column (Cytiva) at 4°C. Sample was eluted at a flow rate of 0.1 ml/min and the elution flowed directly into the x-ray sample cell. The X-ray flux was 7.51 x 10^12^ photons/s with an energy of 11.35 keV and wavelength of 1.094 Å. The sample-detector distance was 1781 mm. Exposures were collected every 1 s throughout the elution until the profile had returned to buffer baseline. The data was analyzed using BioXTAS RAW^65^ **a**nd Guinier and Kratky analysis was performed.

### Autoxidation assays

The *P. azotoformans* sCache_heme protein, *P. putida* protein, or MBP-*P. putida* fusion protein (WT or C59A variant) was degassed overnight in an anaerobic chamber (Coy Laboratory Products) on a 4°C dry bath (Benchmark Scientific). Samples were incubated with a 5x molar excess of potassium ferricyanide for 1 hr to oxidize the heme. Ferricyanide was removed by using a 7k MWCO Zeba spin desalting column (Thermo Scientific). Protein containing Fe^2+^-unligated heme was prepared by incubating with 1 mM dithionite for 1 hr followed by desalting to remove dithionite. To study O_2_ binding, 1.8 mL of 12 µM protein was transferred to a quartz septum-sealed cuvette with a stir bar in an anaerobic chamber and then removed to ambient conditions. UV-Vis spectra were recorded on an Agilent 8543 spectrophotometer for 30 min, stirring at 600 rpm at 20°C. Time traces were initiated by opening the lid of the cuvette and exposing the protein to air. Experiments were done in triplicate. For comparison purposes, rates of oxidation were calculated by fitting the decay in the reduced Soret peak to an exponential decay (y = a^-bx^) after the initial mixing lag time associated with exposing the sample to air.

### Determination of k_off_ for NO complexes

Fe^2+^-unligated protein was prepared as described above. To prepare NO complexes, protein was incubated with a 10x molar excess of nitric oxide donor DEA NONOate (Cayman Chemical). DEA NONOate was then removed by desalting. A CO-dithionite trap was used to measure NO dissociation rate constants. CO was bubbled through degassed buffer containing sodium dithionite. To 1.8 mL of 10 µM protein in an anaerobic septum-sealed cuvette, 200 uL of CO-dithionite solution was injected using a gas-tight syringe to a final dithionite concentration of 30 mM. UV-Vis absorption spectra were recorded over time at 20°C. Dissociation of NO was measured through the decrease in the CO peak (413 nm for MBP-*P. putida* sCache_heme fusion and 415 nm for the *P. azotoformans* FL sCache_heme protein). Experiments were done in triplicate. The change in absorbance was fit to the exponential equation y = a(1 - e^-bt^) + c(1 - e^-dt^) for the MBP-*P. putida* fusion protein and to y = a(1 - e^-bt^) + c for the *P. azotoformans* protein.

### Reactive oxygen species assay

To determine if the proteins were re-oxidizing to the unliganded Fe^3+^ state or the liganded Fe^2+^-O_2_ during the oxygen oxidation experiments, superoxide production was indirectly monitored using the Amplex Red Hydrogen Peroxide/Peroxidase Assay kit (Invitrogen). The kit detects hydrogen peroxide generated by spontaneous dismutation of superoxide to H_2_O_2_. A standard curve was generated using 0-10 μM of H_2_O_2_. The assay was performed in triplicate using 1, 10 and 40 μM of unligated Fe^2+^ protein in a reaction volume of 100 uL in a 96-well plate. The amount of H_2_O_2_ generated due to superoxide release during oxidation on exposing samples to air was hence determined by the reaction of Amplex red with horse radish peroxidase (HRP) via the manufacturer’s specifications.

### Reduction Potential Measurements

Reduction potentials of *P. putida* DUF 3365 WT and C59A variant protein-bound heme cofactors were determined using anaerobic photoreduction with 5-deazariboflavin as previously described 1,2. 29 µM of protein was exchanged into 20 mM sodium phosphate, 15 mM EDTA, 150 mM NaCl, and 20% glycerol (pH 7.0) redox buffer and concentrated to a volume below 100 µL. Samples were rendered anaerobic by incubation in anaerobic glovebox with atmosphere containing 97% N_2_ and 3% H_2_ with ˂ 2ppm O_2_ and kept at 4 °C (Benchmark Scientific MyBlock mini dry bath) for 30 min for oxygen removal. Redox buffer previously deoxygenated in anaerobic glovebox for a minimum of 24 hrs was kept at 4 °C for 30 min in tandem with samples. Following oxygen removal, samples and reagents were transferred to a quartz cuvette with 1 cm pathlength to a final volume of 1.5 mL with redox buffer: 1,2-naphthoquinone redox reference dye and 5-deazoriboflavin were added from a DMSO stock solution to a final concentration of 200 and 5 µM, respectively. UV-visible absorbance spectra were performed in anaerobic glovebox and collected using an Agilent 8453 spectrophotometer maintained at 10 °C with Agilent 89090A temperature controller while stirring. Photochemical reduction was initiated by illuminating the sample with 440 nm Kessil (PR160L-440nm) light at 25% intensity directly 10 cm above cuvette. The sample was illuminated for 10 -13 s and turned off to permit collection of UV-vis spectrum every 15 s; this process was repeated until the end of kinetics reading. Changes in the reduction and oxidation states of the heme cofactor and redox reference dye, respectively, were monitored spectrophotometrically during photoreduction. Absorbance at 550 nm was used to monitor formation of the reduced heme α band and absorbance at 442 and 438 nm for WT and C59A variant, respectively, were used to monitor depletion of oxidized 1,2-naphthoquinone during photoreduction. Data collection continued until absorbance traces reached stable endpoints, indicating complete reduction of the monitored species. The midpoint reduction potential of the protein-bound heme was calculated by comparison with the known reduction potential of 1,2-naphthoquinone at pH 7.0 (E°′ = +145 mV) using the Nernst relation. Nernst terms were calculated using n = 1 for the heme Fe(III)/Fe(II) redox couple and n = 2 for the 1,2-naphthoquinone/1,2-naphthohydroquinone redox couple. For each time point, the ratio of oxidized and reduced species was determined from the absorbance changes between the fully oxidized and fully between the midpoint potential of the reference dye and that of the heme cofactor. The heme midpoint potential was then calculated from the known reduction potential of 1,2-naphthoquinone. Data quality was assessed using several criteria. Formation of the heme α band at 550 nm together with the corresponding shift in the Soret maximum demonstrated retention of protein-bound heme throughout the experiment. Completion of reduction was verified by the presence of stable absorbance plateaus in the kinetic traces. In addition, absorbance changes associated with heme reduction and 1,2-naphthoquinone reduction occurred over overlapping regions of the photoreduction titration trace, indicating that the midpoint potentials of the two redox couples were sufficiently similar for reliable analysis. Finally, Nernst plots were inspected for linear behavior across the reduction range. Datasets yielding Nernst plots exhibiting strong linear correlations (R^2^ > 0.90) were considered acceptable and were used for calculation of midpoint reduction potentials.

### Hemochromagen Assay

Hemochromagen assays were performed according to Bar and Guo. Stock protein solutions were mixed with equal volumes of (0.2 M NaOH, 40% pyridine, 0.5 mM Potassium Ferricyanide), next dithionite was added to 5 mM. The difference between the reduced (550-535 nm) and oxidized (550-535 nm) spectra were used to calculate heme concentrations with extinction coefficients from Berry and Trumpower. Protein concentrations were determined with the BioRad Bradford Assay. Commercial cytochrome c and Hemoglobin from Sigma were used as controls.

NO dissociation rate constants are shown for sCache_heme proteins in comparison to known NO sensors (sGC, H-NOX, NosP) and cytochromes that do not functionally react with NO as part of their functions. Globins, which also bind NO tightly, were excluded because of their varied reactivity with NO that in many cases serves biological roles. Proteins, like mitochondrial cytochrome c, that undergo complex geminate recombination reactions with NO were also excluded.

## Acknowledgements

We thank Kyle Lancaster for supplying the pEC86 plasmid for expression of the cytochrome c maturation factors. This work was supported by NIH grants R35122535 (to B.R.C), R35GM131760 (to I.B.Z.) and Chemical Biology Interface training grant T32GM138826 (to M.D.). CHEXS is supported by the NSF award DMR-2342336, and the MacCHESS resource is supported by NIGMS award 1-P30-GM124166.

## Data Availability

SAXS data was deposited to the small-angle scattering biological data bank (SASDB) under accession code SASD2G3. All data supporting the findings of this study are available within the Article and its Supplementary Information files.

## Author contributions

B.R.C. and I.B.Z. conceived and designed the study. M.D., J.X.,V.M.G., R.D. and H.C. acquired the data. All authors analyzed and interpreted the data. M.D., J.X., B.R.C., and I.B.Z. wrote the manuscript. B.R.C. and I.B.Z. provided resources and supervised the project.

