## Supplementary figures for "A dichotomous family of Cache domain proteins evolved through c-type heme acquisition"

### Supplementary Information: A bacterial nitric-oxide sensor evolved through acquisition of a cytochrome-derived c-type heme-binding motif

Maithili S. Deshpande[1#], Jiawei Xing[2#¶], Vadim M. Gumerov[2], Brian R. Crane[1*], Igor B. Zhulin[2*]

AUTHOR ADDRESS:

[1]Department of Chemistry and Chemical Biology and The Weill Institute for Cell and Molecular Biology, Cornell University, Ithaca, NY 14853 USA

[2]Department of Microbiology and Translational Data Analytics Institute, The Ohio State University, Columbus, OH, 43210 USA

[#]These authors contributed equally to this work. The order of authors was decided alphabetically.

[¶]Current address: Simons Center for Quantitative Biology, Cold Spring Harbor Laboratory, Cold Spring Harbor, NY 11724.

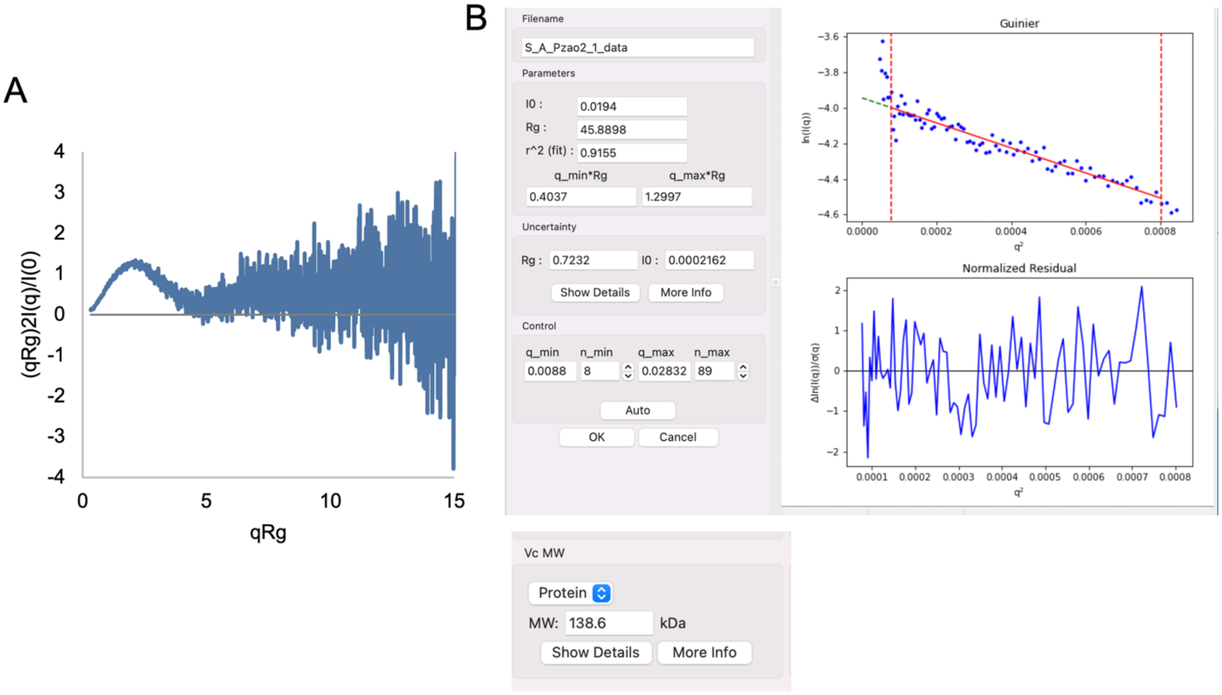

Fig. S1: Size-exclusion chromatography coupled small-angle x-ray scattering (SAXS) of the *P. azotoformans* protein

A. Dimensionless Kratky plot of the *P. azotoformans* dCache domain shows the characteristic Gaussian-shaped peak, indicating that the protein is folded. The peak position is slightly shifted from that of a typical globular protein in which $qR_g = \sqrt{3} \approx 1.73$ and peak height $= 3/e \approx 1.1$.

B. Guinier analysis indicates an $R_g$ of 45.89 and molecular weight generated from volume of correlation (Vc) is 138.6 kDa. The molecular weight of the *P. azotoformans* dimer excluding the detergent micelle is 69 kDa. The molecular weight of a DDM micelle lies between ~70-85 kDa[1,2] thus making the estimated molecular weight of the *P. azotoformans* dimer 139-154 kDa, which is comparable to the molecular weight obtained from the SAXS data.

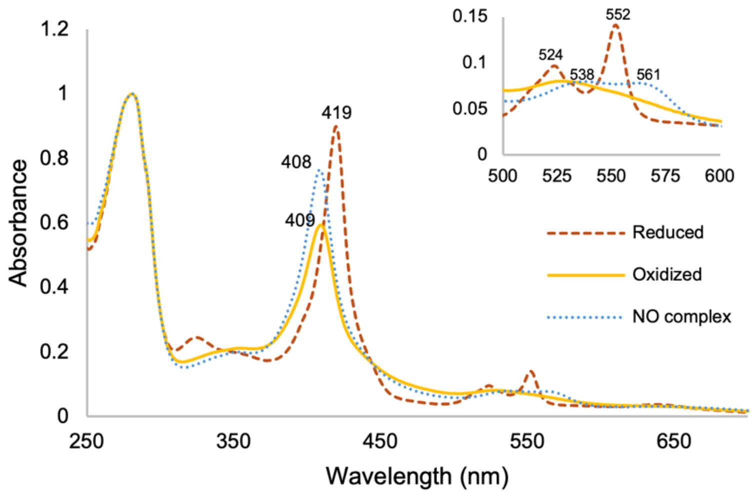

Fig. S2: UV-Vis absorption spectra of the *P. putida* C59A variant

Spectra of the P. putida C59A substitution in oxidized (yellow ━━ ), reduced (red -----) and $Fe^{2+}$-NO forms (blue ·······) with Soret maxima and α/β bands shown. The Soret and α/β bands are shifted from WT, indicating an altered axial ligation state.

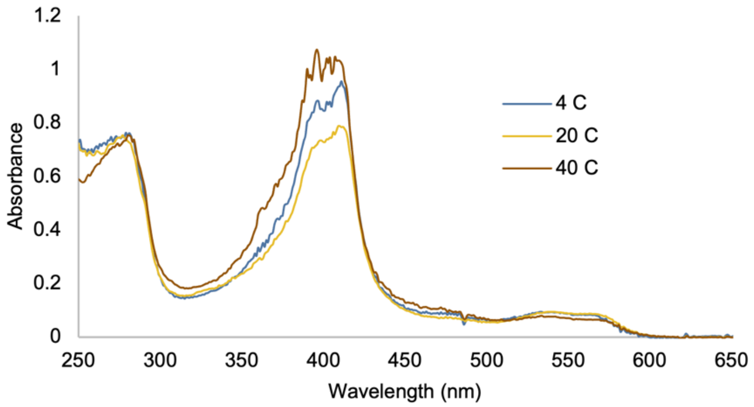

Fig. S3: Temperature-dependent UV-Vis absorption spectra of the $Fe^{2+}$-NO complex of the *P. azotoformans* protein

The mixture of five- and six-coordinate $Fe^{2+}$-NO complexes as seen through absorption at 397 and 412 nm respectively, is maintained at temperatures of 4, 20 and 40 °C.

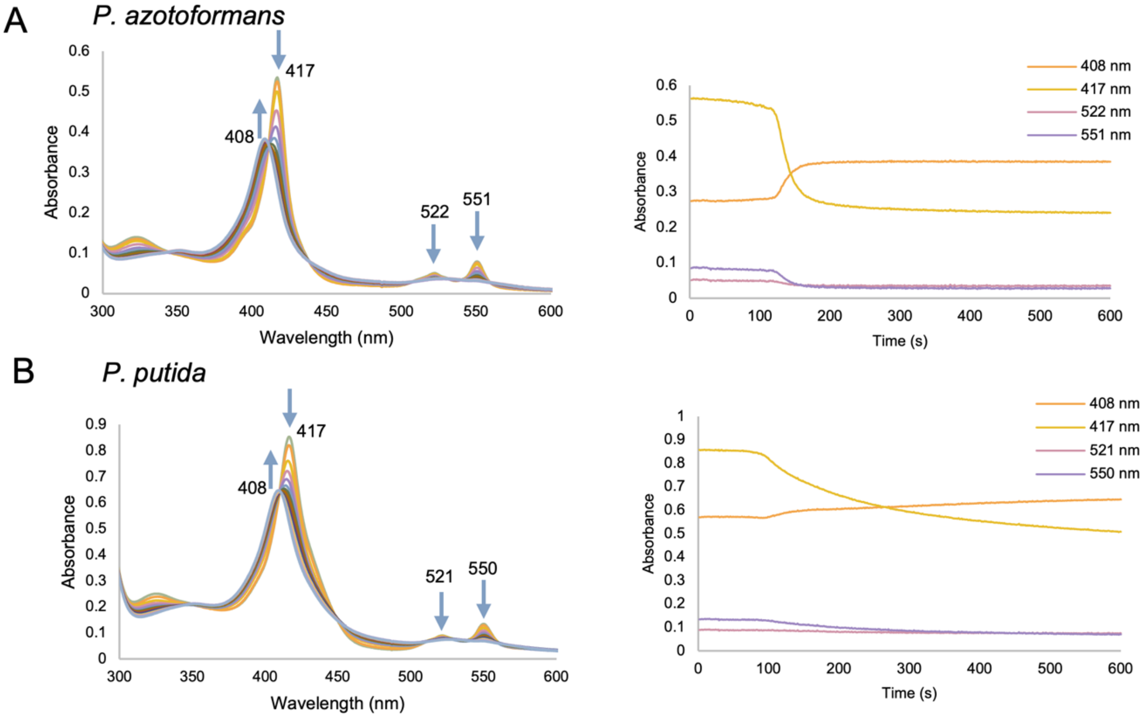

Fig. S4: Auto-oxidation of the *P. azotoformans* and *P. putida* proteins

Upon exposure to air the reduced $Fe^{2+}$ sCache_heme proteins show a decrease in the Soret peak of the reduced species (417 nm) and an increase in the Soret peak of the oxidized species (408 nm), as well as a broadening of the $\alpha$ and $\beta$ peaks. The kinetic traces show the change in different species over the first 600 s. There is an initial lag of ~100 s after which there is an exponential decay in the reduced species. For comparison purposes, apparent re-oxidation rate constants were calculated by fitting the decay in the 417 nm Soret peak to an exponential function ($y = a^{-bx} + c$) after the initial lag time. The *P. azotoformans* protein showed an ~10 x faster rate of oxidation than the *P. putida* protein ($k = (4.78 \pm 0.83) \times 10^{-2}\ s^{-1}$ compared to $k = (0.5 \pm 0.1) \times 10^{-2}\ s^{-1}$).

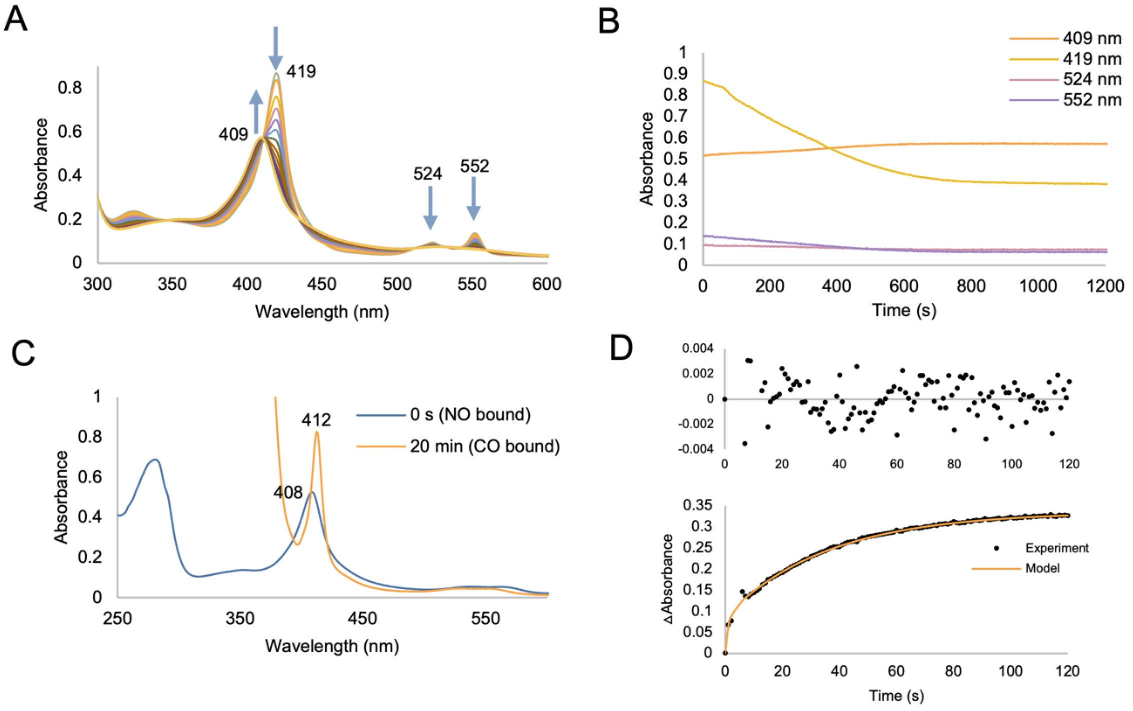

Fig. S5: Auto-oxidation and NO dissociation from *P. putida* C59A

A. The UV-Vis absorption spectra obtained during auto-oxidation of the *P. putida* C59A variant shows an increase in the Soret peak of the oxidized species (409 nm) and a decrease in the Soret peak of the reduced species (419 nm).

B. The time trace shows the change in heme species over 1200 s during auto-oxidation. The heme in this variant is likely destabilized because the decrease in the 419 nm trace in the first 60 s upon removal from the anaerobic chamber in a septum-sealed cuvette even before exposing the protein to air.

C. The UV-Vis absorption spectra of $Fe^{2+}$-NO (blue) and $Fe^{2+}$-CO (orange) during NO dissociation experiments.

D. The fit of the increase in the $Fe^{2+}$-CO absorbance over 120 s during NO dissociation. The fast dissociation indicates that the C59A variant does not bind NO strongly, comparable to WT.

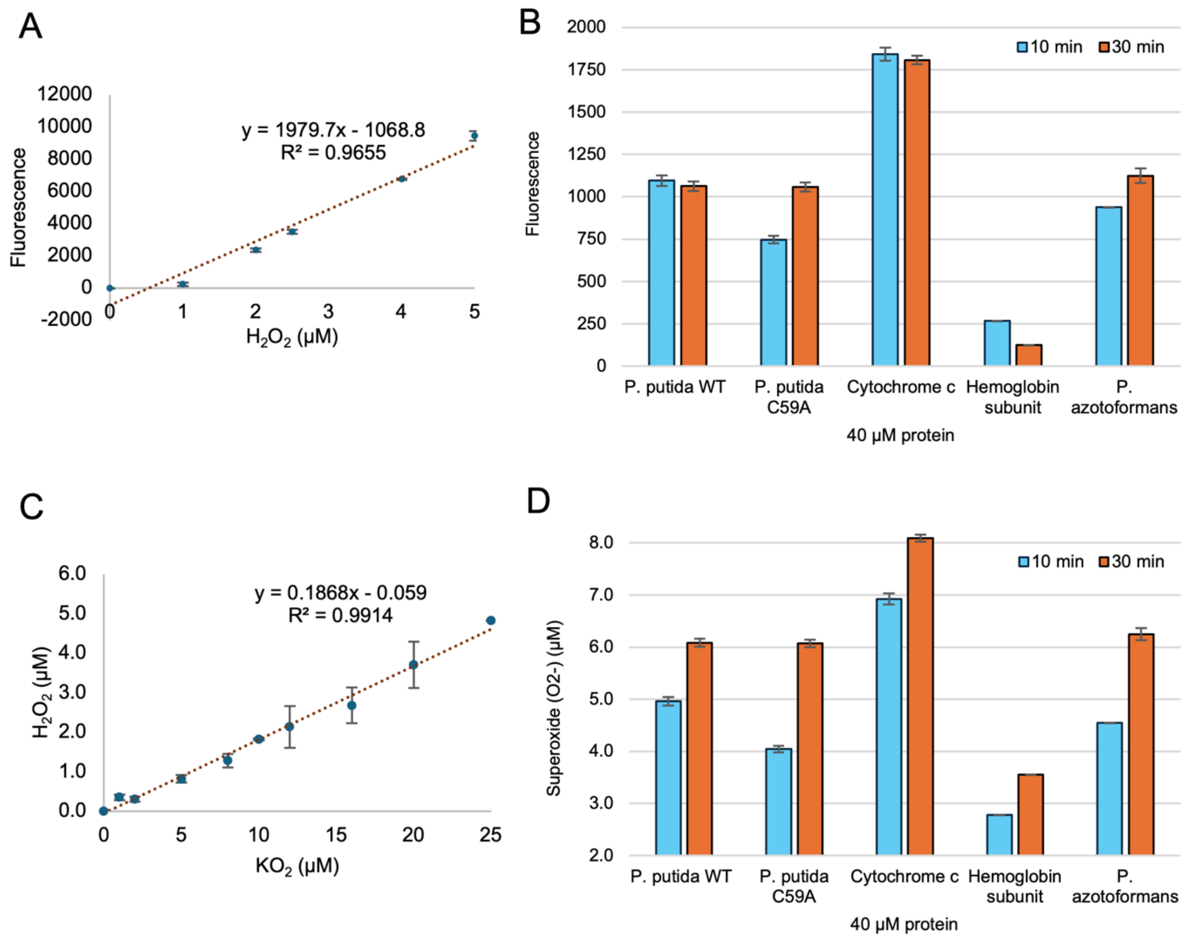

Fig. S6: Release of reactive oxygen species (ROS) from the *P. putida* and *P. azotoformans* proteins upon auto-oxidation

Fluorescence turn-on of Amplex Red on oxidative conversion to resorufin was used to report on ROS release from the sCache_heme proteins during conversion of $Fe^{2+}$ to oxidized $Fe^{3+}$ for 40 μM of the sCache_heme proteins (subunit concentration) compared to cytochrome c ( Cc) and hemoglobin, which respectively produce the $Fe^{3+}$ state or the $Fe^{2+}$-$O_2$ state. Experiments were performed in triplicate.

A. Standard curve generated with $H_2O_2$ concentrations ranging from 0-5 μM at t = 30 min.

B. Fluorescence values at 10 min and 30 min corrected for background fluorescence plotted for 40 μM (subunit concentration) of proteins after auto-oxidation. The $H_2O_2$ produced during oxidation is much lower than the concentration of heme in each sample. This sub-stoichiometric production of $H_2O_2$, which the Amplex Red assay detects, is likely due to the fact that the primary product of auto-oxidation is superoxide, which can disproportionate to $H_2O_2$, but also leads to formation of other ROS. The fluorescence detection of the sCache_heme proteins are 8-9 times higher than hemoglobin at 30 min, indicating that the heme is oxidized to $Fe^{3+}$ and not stably bound as $Fe^{2+}$-$O_2$.

C. Standard curve of the conversion of superoxide (supplied by potassium superoxide) to $H_2O_2$ at t = 30 min using the Amplex Red assay.

D. Superoxide generated by 40 μM (subunit concentration) of protein on autooxidation at 10 min and 30 min. Raw fluorescence values (panel B) were converted to the amount of $H_2O_2$ generated and subsequently related to the amount of superoxide using the standard curve of (C).
